# Deep Unfolded Robust PCA with Application to Clutter Suppression in Ultrasound

**DOI:** 10.1101/469437

**Authors:** Oren Solomon, Regev Cohen, Yi Zhang, Yi Yang, He Qiong, Jianwen Luo, Ruud J.G. van Sloun, Yonina C. Eldar

**Affiliations:** O. Solomon, R. Cohen and Y. C. Eldar are with the Department of Electrical Engineering, Technion-Israel Institute of Technology, Haifa 32000; Y. Zhang is with the department of electrical engineering, Tsinghua University, Beijing 100084, China; Y. Yang, Q. He and J. Luo are with the Department of Biomedical Engineering, Tsinghua University, Beijing 100084; China. R. J. G. van Sloun is with the Department of Electrical Engineering, Eindhoven University of Technology, Eindhoven, The Netherlands

**Keywords:** Ultrasound, Machine learning, Inverse methods, Neural network

## Abstract

Contrast enhanced ultrasound is a radiation-free imaging modality which uses encapsulated gas microbubbles for improved visualization of the vascular bed deep within the tissue. It has recently been used to enable imaging with unprecedented subwavelength spatial resolution by relying on super-resolution techniques. A typical preprocessing step in super-resolution ultrasound is to separate the microbubble signal from the cluttering tissue signal. This step has a crucial impact on the final image quality. Here, we propose a new approach to clutter removal based on robust principle component analysis (PCA) and deep learning. We begin by modeling the acquired contrast enhanced ultrasound signal as a combination of a low rank and sparse components. This model is used in robust PCA and was previously suggested in the context of ultrasound Doppler processing and dynamic magnetic resonance imaging. We then illustrate that an iterative algorithm based on this model exhibits improved separation of microbubble signal from the tissue signal over commonly practiced methods. Next, we apply the concept of deep unfolding to suggest a deep network architecture tailored to our clutter filtering problem which exhibits improved convergence speed and accuracy with respect to its iterative counterpart. We compare the performance of the suggested deep network on both simulations and in-vivo rat brain scans, with a commonly practiced deep-network architecture and the fast iterative shrinkage algorithm, and show that our architecture exhibits better image quality and contrast.

## I. Introduction

MEDICAL ultrasound (US) is a radiation-free imaging modality used extensively for diagnosis in a wide range of clinical segments such as radiology, cardiology, vascular, obstetrics and emergency medicine. Ultrasound-based imaging modalities include brightness, motion, Doppler, harmonic modes, elastography and more [1].

One important imaging modality is contrast-enhanced ultrasound (CEUS) [2] which allows the detection and visualization of blood vessels whose physical parameters such as relative blood volume (rBV), velocity, shape and density are associated with different clinical conditions [3]. CEUS uses encapsulated gas microbubbles as ultrasound contrast agents (UCAs) which are administrated intravenously and are similar in size to red blood cells and thus can flow throughout the vascular system [4]. Among its many applications, CEUS is used for imaging of perfusion at the capillary level [5, 6], for estimating blood velocity in small vessels arteriole by applying Doppler processing [7, 8] and for sub-wavelength vascular imaging [9–14].

A major challenge in ultrasonic vascular imaging such as CEUS is to suppress clutter signals stemming from stationary and slowly moving tissue as they introduce significant artifacts in blood flow imaging [15]. Over the past few decades several approaches have been suggested for clutter removal. The simplest method to remove tissue signal is to filter the ultrasonic signal along the temporal dimension using high-pass finite impulse response (FIR) or infinite impulse response (IIR) filters [16]. However, FIR filters need to have high order while IIR filters exhibit a long settling time which leads to a low number of temporal samples in each spatial location [17] when using focused transmission. The above methods rely on the assumption that tissue motion, if exists, is slow while blood flow is fast. This high-pass filtering approach is prone to failure in the presence of fast tissue motion, as in cardiology, or when imaging microvasculature in which blood velocities are low.

An alternative method for tissue suppression is second harmonic imaging [18], which separates the blood and tissue signals by exploiting the non-linear response of the UCAs to low acoustic pressures, compared with the mostly linear tissue response. This technique, however, limits the frame-rate of the US scanner, and does not remove the tissue signal completely, as tissue can also exhibit a nonlinear response.

The above techniques are based only on temporal information and neglect the high spatial coherence of the tissue, compared to the blood. To take advantage of these spatial characteristics of tissue, a method for clutter removal was presented in [19], based on the singular value decomposition (SVD) of the correlation matrix of successive temporal samples. SVD filtering operates by stacking the (typically beamformed) acquired frames as vectors in a matrix whose column index indicates frame number. Then, an SVD of the matrix is performed and the largest singular values, which correspond to the highly correlated tissue, are zeroed out. Finally, a new matrix is composed based on the remaining singular values and reshaped to produce the blood/UCA movie.

Several SVD-based techniques have been proposed [15, 20–23], such as down-mixing [15] for tissue motion estimation, adaptive clutter rejection for color flow proposed by Lovstakken *et al*. [24] and the principal component analysis (PCA) for blood velocity estimation presented in [25]. However, these methods are based on focused transmission schemes which limit the frame rate and the field of view. This in turn leads to a small number of temporal and spatial samples, reducing the effectiveness of SVD-based filtering. To overcome this limitation, SVD-based clutter removal was extended to ultrafast plane-wave imaging [13, 26–28], demonstrating substantially improved clutter rejection and microvascular extraction. This strategy gained a lot of interest in recent years and nowadays it is used in numerous ultrafast US imaging applications such as functional ultrasound [29, 30], superresolution ultrasound localization microscopy [13, 14] and high-sensitivity microvessel perfusion imaging [26, 27].

A major limitation of SVD-based filtering is the requirement to determine a threshold which discriminates between tissue related and blood related singular values. The appropriate setting of this threshold is typically unclear, especially when the eigenvalue spectra of the tissue and contrast signals overlap. This threshold uncertainty motivates the use of a different model for the acquired data, one that can differentiate between tissue and contrast signals based on the spatio-temporal information, as well as additional information unique to the contrast signal - its sparse distribution in the imaging plane.

Here, we propose two main contributions. The first, is the adaptation of a new model for the tissue/contrast separation problem. We show that similar to other applications such as MRI [31] and recent US Doppler applications [32], we can decompose the acquired, beamformed US movie as a sum of a low-rank matrix (tissue) and a sparse outlier signal (UCAs). This decomposition is also known as robust principle component analysis (RPCA) [33]. We then propose to solve a convex minimization problem to retrieve the UCA signal, which leads to an iterative principal component pursuit (PCP) [33]. Second, we utilize recent ideas from the field of deep learning [34] to dramatically improve the convergence rate and image reconstruction quality of the iterative algorithm. We do so by unfolding [35] the algorithm into a fixed-length deep network which we term Convolutional rObust pRincipal cOmpoNent Analysis (CORONA). This approach harnesses the power of both deep learning and model-based frameworks, and leads to improved performance in various fields [36–40].

CORONA is trained on sets of separated tissue/UCA signals from both *in-vivo* and simulated data. Similar to [37], we utilize convolution layers instead of fully-connected (FC) layers, to exploit the shared spatial information between neighboring image pixels. Our training policy is a two stage process. We start by training the network on simulated data, and then train the resulting network on *in-vivo* data. This hybrid policy allows us to improve the network’s performance and to achieve a fully-automated network, in which all the regularization parameters are also learned. We compare the performance of CORONA with the commonly practiced SVD approach, the iterative RPCA algorithm and an adaptation of the residual network (ResNet), which is considered to be one of the leading deep architectures for a wide variety of problems [41]. We show that CORONA outperforms all other approaches in terms of image quality and contrast.

Unfolding, or unrolling an iterative algorithm, was first suggested by Gregor and LeCun [35] to accelerate algorithm convergence. In the context of deep learning, an important question is what type of network architecture to use. Iterative algorithms provide a natural recurrent architecture, designed to solve a specific problem, such as sparse approximations, channel estimation [42] and more. The authors of [35] showed that by considering each iteration of an iterative algorithm as a layer in a deep network and subsequent concatenation of a few such layers it is possible to train such networks to achieve a dramatic improvement in convergence, i.e., to reduce the number of iterations significantly.

In the context of RPCA, a principled way to construct learnable pursuit architectures for structured sparse and robust low rank models was introduced in [36]. The proposed networks, derived from the iteration of proximal descent algorithms, were shown to faithfully approximate the solution of RPCA while demonstrating several orders of magnitude speed-up compared to standard optimization algorithms. However, this approach is based on a non-convex formulation in which the rank of the low-rank part (or an upper bound on it) is assumed to be known a priori. This poses a network design limitation, as the rank can vary between different applications or even different realizations of the same application, as in CEUS. Thus, for each choice of the rank upper bound, a new network needs to be trained, which can limit its applicability. In contrast, our approach does not require a-priori knowledge of the rank. Moreover, the use of convolutional layers exploits spatial invariance and facilitates our training process as it reduces the number of learnable parameters dramatically.

The rest of the paper is organized as follows. In Section II we introduce the mathematical formulation of the low-rank and sparse decomposition. Section III describes the protocol of the experiments and technical details regarding the realizations of CORONA and ResNet. Section IV presents *in-silico* as well as *in-vivo* results of both the iterative algorithm and the proposed deep networks. Finally, we discuss the results, limitations and further research directions in Section V.

Throughput the paper, *x* represents a scalar, **x** a vector, **X** a matrix and **I**_*N* × *N*_ is the *N* × *N* identity matrix. The notation ||·||_*p*_ represents the standard *p*-norm and ||·||_*F*_ is the Frobenius norm. Subscript *x_l_* denotes the *l*th element of **x** and **x**_*l*_ is the *l*th column of **X**. Superscript **x**^(*p*)^ represents **x** at iteration *p, T*\* denotes the adjoint of **T**, and **Ᾱ** is the complex conjugate of **A**.

## II. Deep Learning Strategy For RPCA In US

### A. Problem formulation

We start by providing a low-rank plus sparse (L+S) model for the acquired US signal. In US imaging, typically a series of pulses are transmitted to the imaged medium. The resulting echoes from the medium are received in each transducer element and then combined in a process called beamforming to produce a focused image. As presented in [43], after demodulation the complex analytical (IQ) signal can be represented as

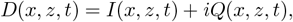

where *I*(*x, z, t*) and *Q*(*x, z, t*) are the in-phase and quadrature components of the demodulated signal, *x, z* are the vertical and axial coordinates, and *t* indicates frame number. The signal *D*(*x, z, t*) is a sum of echoes returned from the blood/CEUS signal *S*(*x, z, t*) as well as from the tissue *L*(*x, z, t*), contaminated by additive noise *N*(*x, z, t*)

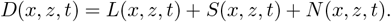

Acquiring a series of movie frames *t* = 1, …, *T*, and stacking them as vectors in a matrix **D**, leads to the following model

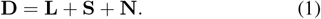

In (1), we assume that the tissue matrix **L** can be described as a low-rank matrix, due to its high spatio-temporal coherence. The CEUS echoes matrix **S** is assumed to be sparse, as blood vessels typically sparsely populate the imaged medium. Assuming that each movie frame is of size *M* × *M* pixels, the matrices in (1) are of size *M*^2^ × *T*. From here on, we consider a more general model, in which the acquired matrix **D** is composed as

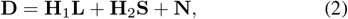

with **H**_1_ and **H**_2_ being the measurement matrices of appropriate dimensions. The model (2) can also be applied to MR imaging, video compression and additional US applications, as we discuss in Section V. Our goal is to formalize a minimization problem to extract both **L** and **S** from **D** under the corresponding assumptions of L+S matrices.

Similar to [31], we propose solving the following minimization problem

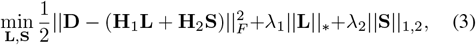

where ||·||_*_ stands for the nuclear norm, which sums the singular values of **L**, and ||·||_1,2_ is the mixed *l*_1,2_ norm, which sums the *l*_2_ norms of each row of **S**. We use the mixed *l*_1,2_ norm since the pattern of the sparse outlier (blood or CEUS signal) is the same between different frames, and ultimately corresponds to the locations of the blood vessels, which are assumed to be fixed, or change very slowly during the acquisition period. The nuclear norm is known to promote low-rank solutions, and is the convex relaxation of the non-convex rank minimization constraint [44].

By defining

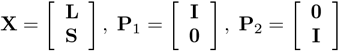

and **A** = [**H**_1_, **H**_2_], (3) can be rewritten as

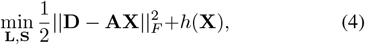

where 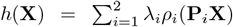 with *ρ*_1_ = ||·||_*_ and *ρ*_2_ = ||·||_1,2_. The minimization problem (4) is a regularized least-squares problem, for which numerous numerical minimization algorithms exist. Specifically, the (fast) iterative shrinkage/thresholding algorithm, (F)ISTA, [45, 46] involves finding the *Moreau’s proximal* (prox) mapping [47, 48] of *h*, defined as

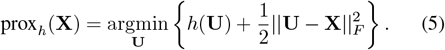

Plugging the definition of **X** into (5) yields

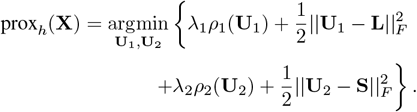

Since prox_*h*_(**X**) is separable in **L** and **S**, it holds that

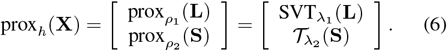

The operators

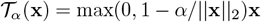

and

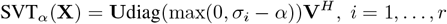

are the mixed *l*_1/2_ soft thresholding [45] and singular value thresholding [49] operators. Here **X** is assumed to have an SVD given by **X** = **UΣV**^*H*^ with **Σ** = diag(*σ_i_*, …, *σ_r_*), a diagonal matrix of the eigenvalues of **X**. The proximal mapping (6) is applied in each iteration to the gradient of the quadratic part of (4), given by

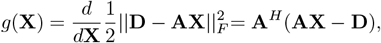

and more specifically,

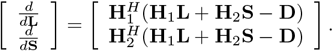

The general iterative step of ISTA applied to minimizing (3) (L+S ISTA) is thus given by

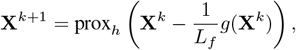

or

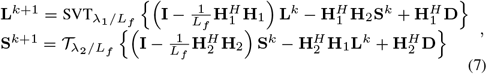

where *L_f_* is the Lipschitz constant of the quadratic term of (4), given by the spectral norm of **A**^*H*^ **A**.

The L+S ISTA algorithm for minimizing (3) is summarized in Algorithm 1. The diagram in Fig. 1(a) presents the iterative algorithm, which relies on knowledge of **H**_1_, **H**_2_ and selection of *λ*_1_ and *λ*_2_.

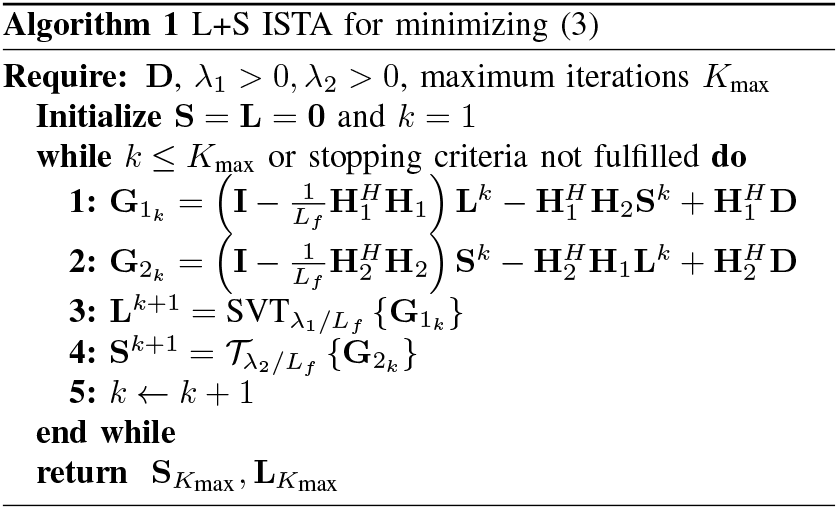

**Fig. 1:**
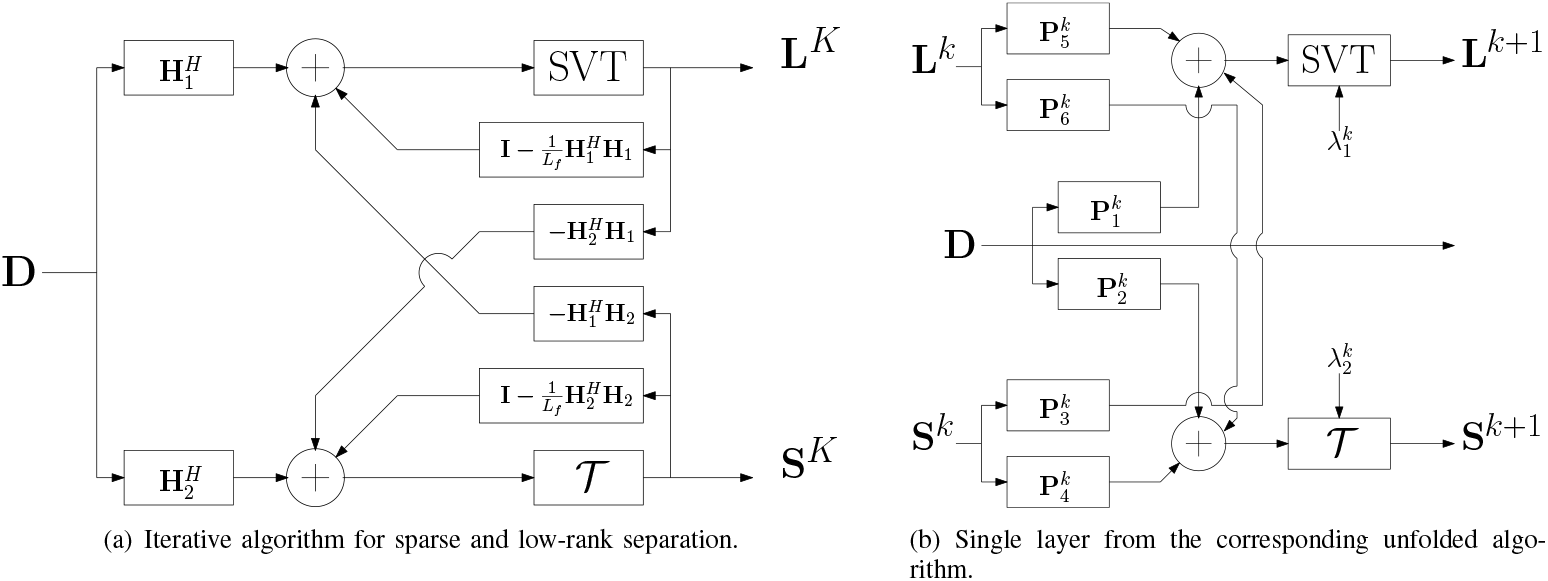
Architecture comparison between the iterative algorithm applied for *K* iterations (panel (a)) and its unfolded counterpart (panel (b)). The learned network in panel (b) draws its architecture from the iterative algorithm, and is trained on examples from a given dataset. In both panels, **D** is the input measurement matrix, and **S**_*k*_ and **L**_*k*_ are the estimated sparse and low-rank matrices in each iteration/layer, respectively.

The dynamic range between returned echoes from the tissue and UCA/blood signal can range from 10dB to 60dB. As this dynamic range expands, more iterations are required to achieve good separation of the signals. This observation motivates the pursuit of a fixed complexity algorithm. In the next section we propose CORONA which is based on unfolding Algorithm 1. Background on learning fast sparse approximations is given in Section I of the supplementary materials.

### B. Unfolding the iterative algorithm

An iterative algorithm can be considered as a recurrent neural network, in which the *k*th iteration is regarded as the *k*th layer in a feedforward network [36]. To form a convolutional network, one may consider convolutional layers instead of matrix multiplications. With this philosophy, we form a network from (7) by replacing each of the matrices dependent on **H**_1_ and **H**_2_ with convolution layers (kernels) 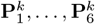 of appropriate sizes. These will be learned from training data. Contrary to previous works in unfolding RPCA which considered training fully connected (FC) layers [36], we employ convolution kernels in our implementation which allows us to achieve spatial invariance while reducing the number of learned parameters considerably.

The kernels as well as the regularization parameters 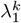 and 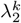 are learned during training. By doing so, the following equations for the *k*th layer are obtained

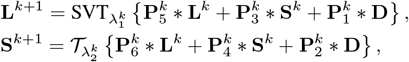

with ∗ being a convolution operator. The latter can be considered as a single layer in a multi-layer feedforward network, which we refer to as CORONA: Convolutional rObust pRincipal cOmpoNent Analysis. A diagram of a single layer from the unfolded architecture is given in Fig. 1(b), where the supposedly known model matrices were replaced by the 2D convolution kernels 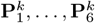, which are learned as part of the training process of the overall network.

In many applications, the recovered matrices **S** and **L** represent a 3D volume, or movie, of dynamic objects imposed on a (quasi) static background. Each column in **S** and **L** is a vectorized frame from the recovered sparse and low-rank movies. Thus, we consider in practice our data as a 3D volume and apply 2D convolutions. The SVT operation (which has similar complexity as the SVD operation) at the *k*th layer is performed after reshaping the input 3D volume into a 2D matrix, by vectorizing and column-wise stacking each frame.

The thresholding coefficients are learned independently for each layer. Given the *k*th layer, the actual thresholding values for both the SVT and soft-thresholding operations are given by 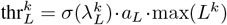 and 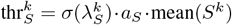 respectively, where *σ*(*x*) = 1/(1 + exp(−*x*)) is a sigmoid function, *a_L_* and *a_S_* are fixed scalars (in our application we chose *a_L_* = 0.4 and *a_S_* = 1.8) and 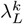 and 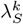 are learned in each layer by the network.

### C. Training CORONA

CORONA is trained using back-propagation in a supervised manner. Generally speaking, we obtain training examples **D**_*i*_ and corresponding sparse **Ŝ**_*i*_ and low-rank 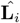 decompositions. In practice, **Ŝ**_*i*_ and 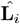 can either be obtained from simulations or by decomposing **D**_*i*_ using iterative algorithms such as FISTA [50]. The loss function is chosen as the sum of the mean squared errors (MSE) between the predicted **S** and **L** values of the network and **Ŝ**_*i*_, 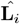, respectively,

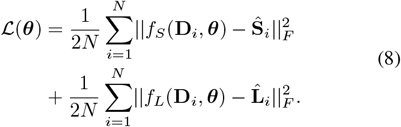

In the latter equation, *f_S_/f_L_* is the sparse/low-rank output of CORONA with learnable parameters 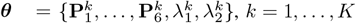, where *K* is the number of chosen layers.

Training a deep network typically requires a large amount of training examples, and in practice, US scans of specific organs are not available in abundance. To be able to train CORONA, we thus rely on two strategies: patch-based analysis and simulations. Instead of training the network over entire scans, we divide the US movie used for training into 3D patches (axial coordinate, lateral coordinate and frame number). Then we apply Algorithm 1 on each of these 3D patches. The SVD operations in Algorithm 1 become computationally tractable since we work on relatively small patches. The resulting separated UCA movie is then considered as the desirable outcome of the network and the network is trained over these pairs of extracted 3D patches from the acquired movie, and the resulting reconstructed UCA movies. In practice, the CEUS movie used for training is divided into 3D patches of size 32 × 32 × 20 (32 × 32 pixels over 20 consecutive frames) with 50% overlap between neighboring patches. The regularization parameters of Algorithm 1, *λ*_1_ and *λ*_2_ are chosen empirically, but are chosen once for all the extracted patches.

In Section VI of the supplementary materials, we provide a detailed description of how the simulations of the UCA and tissue movies were generated. In particular, we detail how individual UCAs were modeled and propagated in the imaging plane, and describe the cluttering tissue signal model. We then demonstrate the importance of training on both simulations and *in-vivo* data in Section IV of the supplementary materials.

## III. Experiments

The brains of two rats were scanned using a Vantage 256 system (Verasonics Inc., Kirkland, WA, USA). An L20-10 probe was utilized, with a central frequency of 15MHz. The rats underwent craniotomy after anesthesia to obtain an imaging window of 6 × 2mm^2^. A bolus of 100*μ*L SonoVue™ (Bracco, Milan, Italy) contrast agent, diluted with normal saline with a ratio of 1:4, was administered intravenously to the rats tail vein. Plane-wave compounding of five steering angles (from −12° to 12°, with a step of 6°) was adopted for ultrasound imaging. For each rat, over 6000 consecutive frames were acquired with a frame rate of 100Hz. 300 frames with relatively high B-mode intensity were manually selected for data processing in this work.

In recent years, several deep learning based architectures have been proposed and applied successfully to classification problems. One such approach is the residual network, or ResNet [41]. ResNet utilizes convolution layers, along with batch normalization and skip connections, which allow the network to avoid vanishing gradients and reduce the overall number of network parameters.

To compare with CORONA, we implemented ResNet using complex convolutions for the tissue clutter suppression task. The network does not recover the tissue signal, as CORONA, but only the UCA signal. In Section IV and in the supporting materials file, we compare both architectures and assess the advantages and disadvantages of each network. In Section IV-B, we show that CORONA outperforms ResNet in terms of image quality (contrast) of the CEUS signal.

Both ResNet and CORONA were implemented in Python 3.5.2, using the PyTorch 0.4.1 package. CORONA consists of 10 layers. First three layers used convolution filters of size 5 × 5 × 1 with stride (1, 1, 1), padding (2, 2, 0) and bias, while the last seven layers used filters of size 3 × 3 × 1 with stride (1, 1, 1), padding (1, 1, 0) and bias. Training was performed using the ADAM optimizer with a learning rate of 0.002. For the *in-vivo* experiments in Section IV, we trained the network over 2400 simulated training pairs and additional 2400 *in-vivo* pairs taken only from the first rat. Training pairs were generated from the acquired US clips, after dividing each clip to 32 × 32 × 20 patches. We then applied Algorithm 1 for each patch with *λ*_1_ = 0.02, *λ*_2_ = 0.001 and *D*_max_ = 30000 iterations to obtain the separated UCA signal for the training process. Algorithm 1 was implemented in MATLAB (Mathworks Inc.) and was applied to the complex-valued IQ signal. PyTorch performs automatic differentiation and back-propagation using the Autograd functionality, and version 0.4.1 also supports back-propagation through SVD, but only for real valued numbers. Thus, complex valued convolution layers and SVD operations were implemented.

## IV. RESULTS

### A. Simulation results

In this section we provide reconstruction results for CORONA applied to a simulated dataset, and trained on simulations. Figure 2 presents reconstruction results of the UCA signal **S** and the low-rank tissue **L** against the ground truth images. Panel (a) shows a representative image in the form of maximum intensity projection (MIP)^1^ of the input cluttered movie (50 frames). It is evident that the UCA signal, depicted as randomly twisting lines, is masked considerably by the simulated tissue signal. Panel (b) illustrates the ground truth MIP image of the UCA signal, while panel (c) presents the MIP image of the recovered UCA signal via CORONA. Panels (d) and (e) show MIP images of the ground truth and CORONA recovery, respectively.

**Fig. 2:**
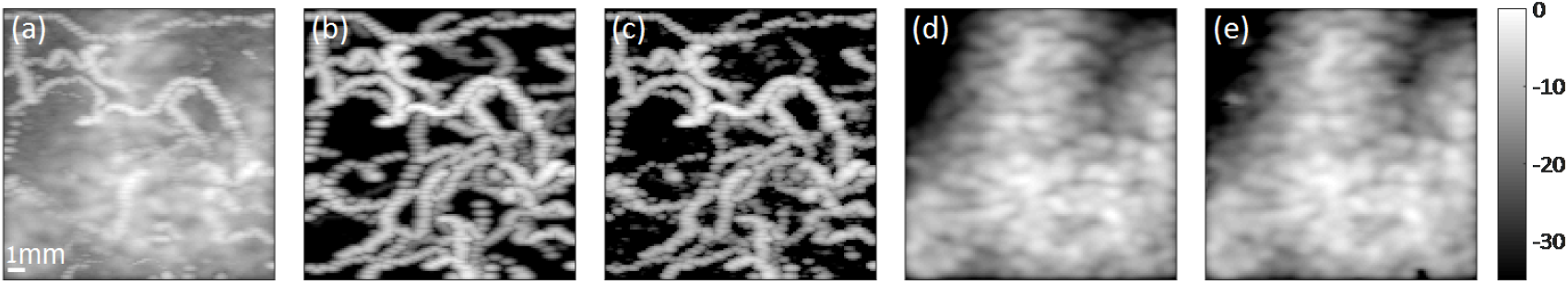
Simulation results of CORONA. (a) MIP image of the input movie, composed from 50 frames of simulated UCAs cluttered by tissue. (b) Ground-truth UCA MIP image. (c) Recovered UCA MIP image via CORONA. (d) Ground-truth tissue MIP image. (d) Recovered tissue MIP image via CORONA. Color bar is in dB.

Observing all panels, it is clear that CORONA is able to recover reliably both the UCA signal and the tissue signal. Section II in the supporting materials provides additional simulation results, showing also the recovered UCA signal by ResNet. Although qualitatively ResNet manages to recover well the UCA signal, its contrast is lower than the contrast of the CORONA recovery, which presents a clearer depiction of the random vascular structure of the simulation. Moreover, ResNet does not recover the tissue signal, while CORONA does.

As CORONA draws its architecture from the iterative ISTA algorithm, our second aim in this section is to assess the performance of both CORONA and the FISTA algorithm by calculating the MSE of each method as a function of iteration/layer number. Each layer in CORONA can be thought of as an iteration in the iterative algorithm. To that end, we next quantify the MSE over the simulated validation batch (sequence of 100 frames) as a function of layer number (CORONA) and iteration number (FISTA), as presented in Fig. 3. For both methods, the MSE for the recovered sparse part (UCA signal) **S** and the low-rank part (tissue signal) **L** were calculated as a function of iteration/layer number, as well as the average MSE of both parts, according to (8) (*α* = 0.5). For each layer number, we constructed an unfolded network with that number of layers, and trained it for 50 epochs on simulated data only.

**Fig. 3:**
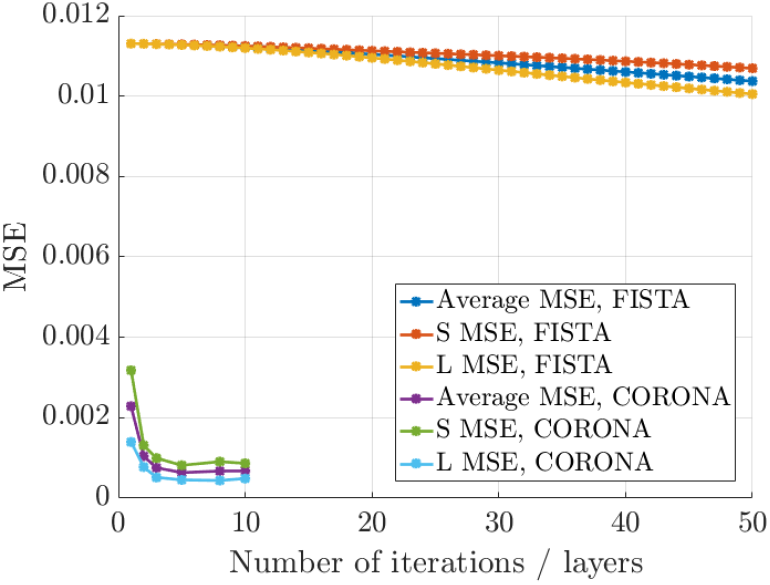
MSE plot for the FISTA algorithm and CORONA as a function of the number of iterations/layers.

Observing Fig. 3, it is clear that even when considering CORONA with only 1 layer, its performance in terms of MSE in an order of magnitude better than FISTA applied with 50 iterations. Adding more layers improves the CORONA MSE, though after 5 layers, the performance remains roughly the same. Figure 3 also shows that a clear decreasing trend is present for the FISTA MSE, however a dramatic increase in the number of iterations is required by FISTA to achieve the same MSE values.

### B. In-vivo experiments

We now proceed to demonstrate the performance of CORONA on *in-vivo* data. As was described in Section III, CORONA was trained on both simulated and experimental data. In Fig. 4, panel (a) depicts SVD based separation of the CEUS signal, panel (b) shows the FISTA based separation and panel (c) shows the result of CORONA. The lower panels of Fig. 4 also compare the performance of the trained ResNet (panel (f)) on the *in-vivo* data as well as provide additional comparison to the commonly used wall filtering. Specifically, we use a 6th order Butterworth filter with two cutoff frequencies of 0.2*π* (panel (d)) and 0.9*π* (panel (e)) radians/samples. Two frequencies were chosen which represent two scenarios. The cutoff frequency of the recovery in panel (d) was chosen to suppress as much tissue signal as possible, without rejecting slow moving UCAs. In panel (e), a higher frequency was chosen, to suppress the slow moving tissue signal even further, but as can be seen, at a cost of removing also some of the slower bubbles. The result is a less consistent vascular image. Visually judging, all panels of Fig. 4 shows that ResNet outperforms both the SVD and wall filtering approaches. However, a more careful observation shows that the ResNet output, although more similar to CORONA’s output, seems more grainy and less smooth than CORONA’s image. CORONA’s recovery exhibits the highest contrast, and produces the best visual.

**Fig. 4:**
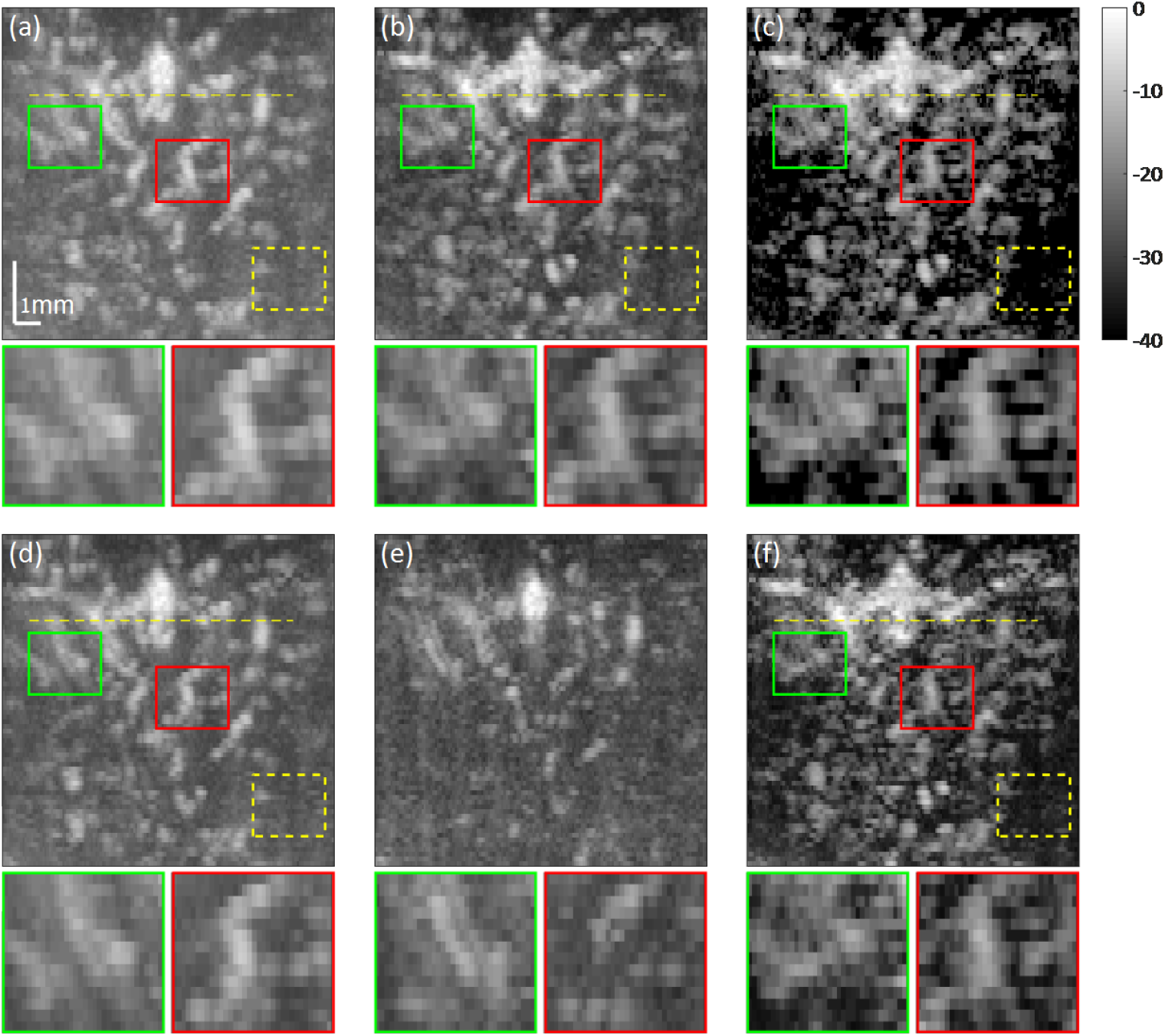
Recovery of *in-vivo* CEUS signal depicting rat brain vasculature. (a) SVD based separation. (b) L+S FISTA separation. (c) Deep network separation, with the unfolded architecture of the FISTA algorithm. (d) Wall filtering with cutoff frequency of 0.2*π* (e) Wall filtering with cutoff frequency of 0.9*π* (f) ResNet. Color bar is in dB.

In each panel, the green and red boxes indicate selected areas, whose enlarged views are presented in the corresponding green and red boxes below each panel. Visual inspection of the panels (a)-(f) shows that FISTA, ResNet and CORONA achieve CEUS signal separation which is less noisy than the naive SVD approach and wall filtering. Considering the enlarged regions below the panels further supports this conclusion, showing better contrast of the FISTA and deep networks outputs. The enlarged panels below panels (d) and (e) show that indeed, as the cutoff frequency of the wall filter is increased, slow moving UCAs are also filtered out. Both deep networks exhibit higher contrast than the other approaches.

To further quantify the performance of each method, we provide two metrics to assess the contrast ratio of their outputs, termed contrast to noise ratio (CNR) and contrast ratio (CR).

CNR is calculated between a selected patch, e.g. the red or green boxes in panels (a)-(f) and a reference patch, marked by the dashed yellow patches, which represents the background, for the same image. That is, for each panel we estimate the CNR values of the red - yellow and green - yellow boxes, where *μ_s_* is the mean of the red/green box with variance 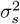 and *μ_b_* is the mean of the dashed yellow patch with variance 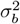. The CNR is defined as

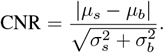

Similarly, the CR is defined as

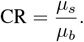

Table I and Table II provide the calculated CNR and CR values of each method, respectively.

**TABLE I:**
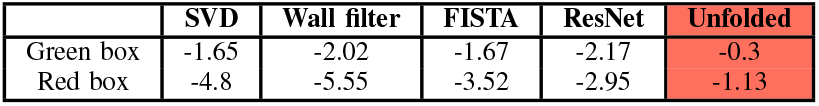
CNR values for the selected green and red rectangles of Fig. 4, as compared with the dashed yellow background rectangle in each corresponding panel. All values are in dB.

**TABLE II:** CR values for the selected green and red rectangles of Fig. 4, as compared with the dashed yellow background rectangle in each corresponding panel. All values are in dB.

|  | SVD | Wall filter | FISTA | ResNet | Unfolded |
| --- | --- | --- | --- | --- | --- |
| Green box | 4.68 | 4.5 | 5.52 | 7.92 | 15.24 |
| Red box | 4.56 | 4.1 | 5.24 | 7.55 | 14.88 |

In both metrics, higher values imply higher contrast ratios, which suggest better noise suppression and better signal depiction. Considering both tables, CORONA outperforms all other approaches. In most cases, its performance is an order of magnitude better than that SVD. The CR values of ResNet are also better than the baseline SVD, though lower than those of CORONA. Its CNR values however, are not always higher than those of the SVD. In terms of CR, the FISTA results are better than those of the SVD filter, though lower than the deep-learning based approaches. In terms of CNR, for the green box, FISTA is comparable to SVD and better than ResNet, while for the red box, its performance is the worst.

Both metrics support the previous conclusions, that by combining a proper model to the separation problem with a data-driven approach leads to improved separation of UCA and tissue signals, as well as noise reduction as compared to the popular SVD approach.

Finally, we also provide intensity cross-sections, taken along the horizontal yellow dashed line for each method, as presented in Fig. 5. Considering the intensity cross-section of Fig. 5, it is evident that all methods reconstruct the peaks with good correspondence. The FISTA and deep-learning networks’ profiles exhibit higher contrast than the SVD and wall filter (deeper “cavities”). In some areas, the unfolded (yellow) profiles seems to vanish. This is because the attained value is −∞. The supporting materials file contains additional comparisons. Section III presents the training and validation losses of the networks, as well as the evolution of the regularization coefficients of CORONA as a function of epoch number. Section IV discusses the importance of training the networks on both simulations and *in-vivo data* when applying CORONA on *in-vivo* experiments, while Section V presents the training and execution times for both networks.

**Fig. 5:**
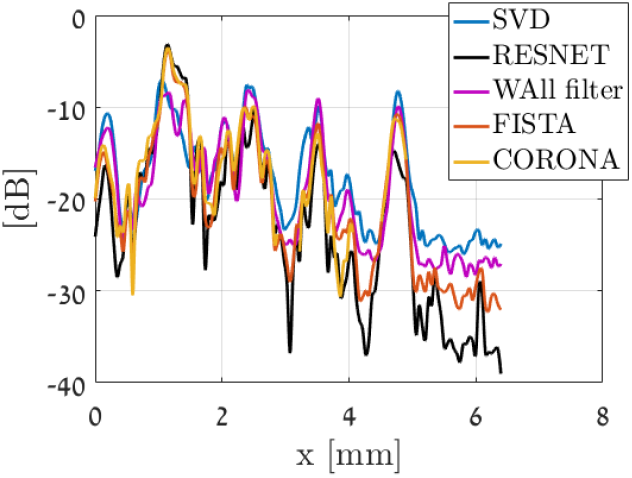
Intensity profiles across the dashed yellow lines in panels (a)-(c) of Fig. 4. Regions in which CORONAs’ curve is missing indicate a value of −∞. Values are in dB.

## V. DISCUSSION AND CONCLUSIONS

In this work, we proposed a low-rank plus sparse model for tissue/UCA signal separation, which exploits both spatio-temporal relations in the data, as well as the sparse nature of the UCA signal. This model leads to a solution in the form of an iterative algorithm, which outperforms the commonly practiced SVD approach. We further suggested to improve both execution time and reconstructed image quality by unfolding the iterative algorithm into a deep network, referred to as CORONA. The proposed architecture utilizes convolution layers instead of FC layers and a hybrid simulation-*in-vivo* training policy. Combined, these techniques allow CORONA to achieve improved performance over its iterative counterpart, as well as over other popular architectures, such as ResNet. We demonstrated the performance of all methods on both simulated and *in-vivo* datasets, showing improved vascular depiction in a rat’s brain.

We conclude by discussing several points, regarding the performance and design of deep-learning based networks. First, we attribute the improved performance over the commonly practiced SVD filtering, wall filtering and FISTA to two main reasons. The first, is the fact that for application on *in-vivo* data, the networks are trained based on both *in-vivo* data and simulated data. The simulated data provides the networks with an opportunity to learn from “perfect” examples, without noise and with absolute separation of UCAs and their surroundings. In Section IV of the supplementary materials we show the effect on recovery when the network is trained with and without experimental data. The iterative algorithm, on the other hand, cannot learn or improve its performance on the *in-vivo* data from the simulated data. The second, is the fact that both networks rely on 2D complex convolutions. Contrary to FC layers, convolution layers reduce the number of learnable parameters considerably, thus help avoid over-fitting and achieve good performance even when the training sets are relatively low. Moreover, convolutions offer spatial invariance, which allows the network to capture spatially translated UCAs.

Focusing on patch-based training (Section II-C) over entire image training has several benefits. UCAs are used to image blood vessels, and as such entire images will include implicitly blood vessel structure. Thus, training over entire images may result in the network being biased towards the vessel trees presented in the (relatively small) training cohort. On the other hand, small patches are less likely to include meaningful structure, hence training on small patches will be less likely to bias the network towards specific blood vessel structures and enable the network to generalize better. Furthermore, as FISTA and CORONA employ SVD operations, processing the data in small batches improves execution time [27, 51].

Second, as was mentioned in the introduction, in the context of RPCA, a principled way to construct learnable pursuit architectures for structured sparse and robust low rank models was introduced in [36]. The proposed network was shown to faithfully approximate the RPCA solution with several orders of magnitude speed-up compared to its standard optimization algorithm counterpart. However, this approach is based on a non-convex formulation of the nuclear norm in which the rank (or an upper bound of it) is assumed to be known a priori.

The main idea in [36] is to majorize the non-differentiable nuclear norm with a differentiable term, such that the low-rank matrix is factorized as a product of two matrices, **L** = **AB**, where **A** ∈ ℝ^*n*×*q*^ and **B** ∈ ℝ^*q*×*m*^. Using this kind of factorization alleviates the need to compute the SVD product, but introduces another unknown parameter *q* which needs to be set (typically by hand), and corresponds to the rank of the low-rank matrix. This poses a network design limitation, as the rank can vary between different applications or even different realizations of the same application, requiring the network to be re-trained per each new choice of *q*.

In fact, this is the same rank-thresholding parameter as in the standard SVD filtering technique, which we want to avoid hand-tuning. Moreover, this kind of factorization leads to a non-convex minimization problem, whose globally optimal stationary points depend on the choice of the regularization parameter *λ*_*_. Since typically these parameters are chosen empirically, a wrong choice of *λ*_*_ may lead to suboptimal reconstruction results of the RPCA problem, which are then used as training data for the fixed complexity learned algorithm. Since we operate on the original convex problem, we train against optimal reconstruction results of the RPCA algorithm, without the need to a-priori estimate the low-rank degree, *q*.

Third, currently CORONA and ResNet offer a trade-off between them. By relying on convolutions, CORONA is trained with a considerable lower number of parameters (314 for 1 layer, 1796 for 10 layers) than the ResNet (25378). CORONA outperforms ResNet in both visual quality and quantifiable metrics, as presented in Section IV. However, its training and execution times are slower (see Section V in the supporting materials file). This performance-runtime trade-off is attributed to the fact that CORONA relies on SVD decomposition in each layer, which is a relatively computationally demanding operation. However, it allows the network to learn the rank of the low-rank matrix, without the need to upper bound it and restrict the architecture of the network. Incorporation of fast approximations for SVD computations, such as truncated or random SVD [51–54], can potentially expedite the network’s performance and achieve faster execution than ResNet. It is also important to keep in mind that ResNet does not recover the tissue signal, only the UCA signal. In some applications, such as super-resolution CEUS imaging over long time durations, the tissue signal is used to correct for motion artifacts.

On a final note, the proposed iterative and deep methods were demonstrated on the extraction of CEUS signal from an acquired IQ movie, but in principle can also be applied to dynamic MRI sequences, as well as to the separation of blood from tissue, e.g. for Doppler processing. In the latter case, the dynamic range between the tissue signal and the blood signal will be greater than that of the tissue and UCA signal. In terms of the iterative algorithm, this would lead to more iterations for the separation process, but once the iterative algorithm has finished, its learned version could be trained on its output to achieve faster execution.

## Supporting information

## Acknowledgment

The authors would like to thank De Ma and Zhifei Dai from the Biomedical Engineering department of Peking university for help in performing the *in-vivo* experiments.

## Footnotes

1 In order to present a single representative image, we take the pixel-wise maximum from each movie. This process is also referred to as maximum intensity projection, and is a common method to visualize CEUS images.

