## Supplementary material for "Deep Unfolded Robust PCA with Application to Clutter Suppression in Ultrasound"

### I. Learning fast approximations via unfolding

To better understand the concept of unfolding an iterative algorithm, we briefly describe the basic ideas presented in [1]. Consider the following sparse recovery model

$$\mathbf{y} = \mathbf{A}\mathbf{x},$$

where $\mathbf{y}$ is a length-$m$ measurement vector, $\mathbf{x}$ is a length-$n$ sparse vector to be recovered, and $\mathbf{A}$ is the

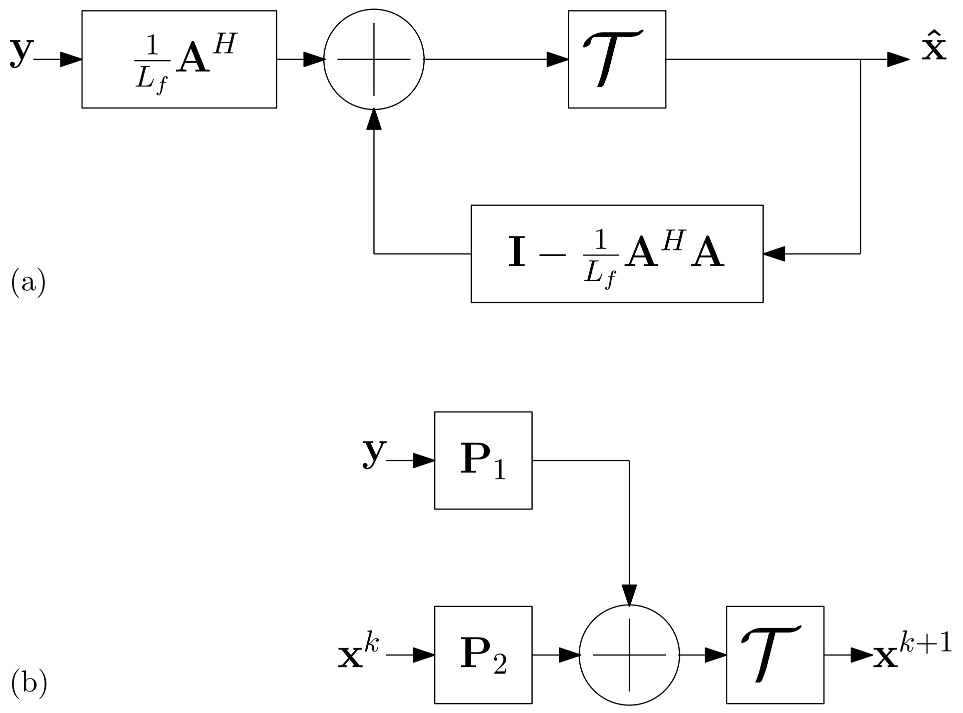


Figure 1: ISTA iterative algorithm (panel (a)) compared with the learned ISTA (panel (b)). Each iteration in the iterative algorithm is replaced with a single layer in the learned algorithm. Instead of using the model parameters such as $\mathbf{A}$, these parameters are replaced with general matrices $\mathbf{P}_1$ and $\mathbf{P}_2$ which are learned.

sensing matrix. Recovering $\mathbf{x}$ from $\mathbf{y}$ can be performed by formulating the following convex minimization problem

$$\min_{\mathbf{x}} \frac{1}{2}||\mathbf{y}-\mathbf{A}\mathbf{x}||_2^2+\lambda||\mathbf{x}||_1, \tag{1}$$

where $\lambda > 0$ is a regularization parameter. A popular iterative algorithm which minimizes (1) is the ISTA algorithm, or its faster counterpart, the fast ISTA (FISTA). FISTA is guaranteed to converge, in the worst case scenario, with a rate proportional to $1/k^2$, with $k$ being the iteration number. As suggested in [1], this convergence can be sped up by proposing a learned version of ISTA (LISTA). Furthermore, the authors of [2] demonstrated that the unfolded architecture facilitates a trade-off between fast convergence and reconstruction accuracy of the sparse recovery problem.

More specifically, the iterative scheme of ISTA consists of the following iterative step

$$\mathbf{x}^{k+1}=\mathcal{T}_{\lambda/L_f}\left\{\left(\mathbf{I}-\frac{1}{L_f}\mathbf{A}^H\mathbf{A}\right)\mathbf{x}^k+\frac{1}{L_f}\mathbf{A}^H\mathbf{y}\right\},$$

with $\mathcal{T}_{\lambda/L_f}(\cdot)$ being the element-wise soft-thresholding operator with parameter $\lambda/L_f$ and $L_f$ is the spectral norm of $\mathbf{A}^H\mathbf{A}$. This iterative procedure is illustrated in panel (a) of Fig. 1, where $\hat{\mathbf{x}}$ is the output of the ISTA algorithm.

Conversely, we can consider each iteration of the iterative algorithm in panel (a) of Fig. 1 as a single layer in a feedforward network. Instead of using the known matrix $\mathbf{A}$ we replace the matrices in panel (a) with general matrices $\mathbf{P}_1$ and $\mathbf{P}_2$ to be learned, as well as the regularization parameter $\lambda$, as illustrated in panel (b) of Fig. 1. Thus, a single layer of this unfolded network is described by

$$\mathbf{x}^{k+1}=\mathcal{T}_{\lambda/L_f}\left\{\mathbf{P}_2\mathbf{x}^k+\mathbf{P}_1\mathbf{y}\right\}.$$

By concatenating several such layers (typically less than ten layers, corresponding to ten iterations are sufficient), a deep network is formed.

### II. ResNet architecture

In this section we provide an additional result of ResNet applied to the simulated data (trained for 10 epochs on simulated data), as well as a detailed description of the complex ResNet architecture.

Figure 2 presents the ResNet recovery of the same simulated movie presented in Fig. 2 of the main paper. Visual inspection of Fig. 2 reveals that in the case of simulations, ResNet suppresses the tissue signal and reveals the UCA signal. However, comparing panel (c) of Fig. 2 to panel (c) of Fig. 2 of the main paper shows that the CORONA reconstruction achieves higher contrast, in line with the conclusions drawn in the main paper. Moreover, CORONA is able to recover the tissue signal as well as the UCA signal, whereas ResNet recovers the UCA signal only. Figure 3 shows the ResNet architecture used in

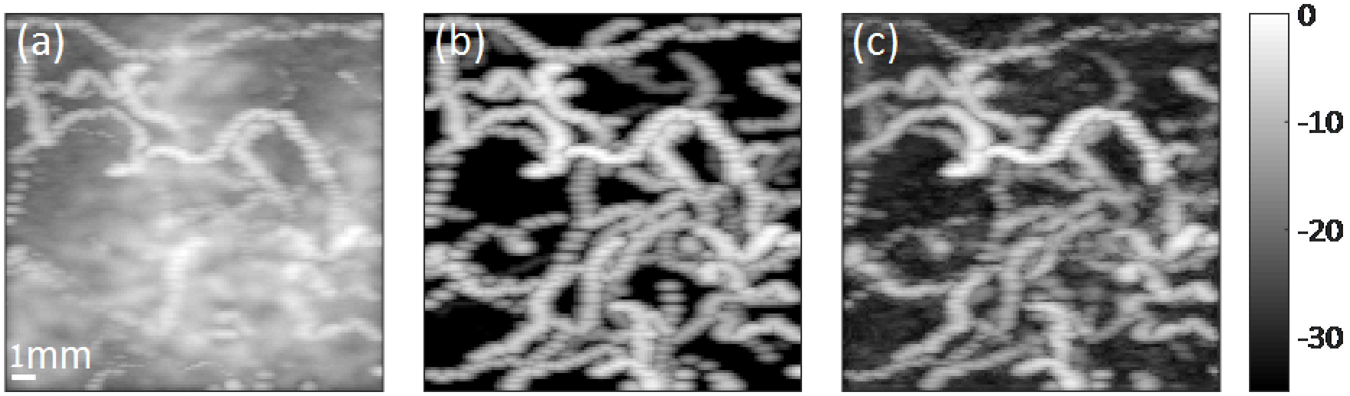


Figure 2: Simulation results of ResNet. (a) MIP image of the same simulated movie used in the main paper. (b) Ground truth MIP image of the UCAs. (c) MIP image of the UCAs recovered by ResNet. Color bar is in dB.

this work. Here, Conv. layer is a complex convolution layer, and $16@5 \times 5$ refers to 16 convolution channels with a $5 \times 5$ pixels kernel.

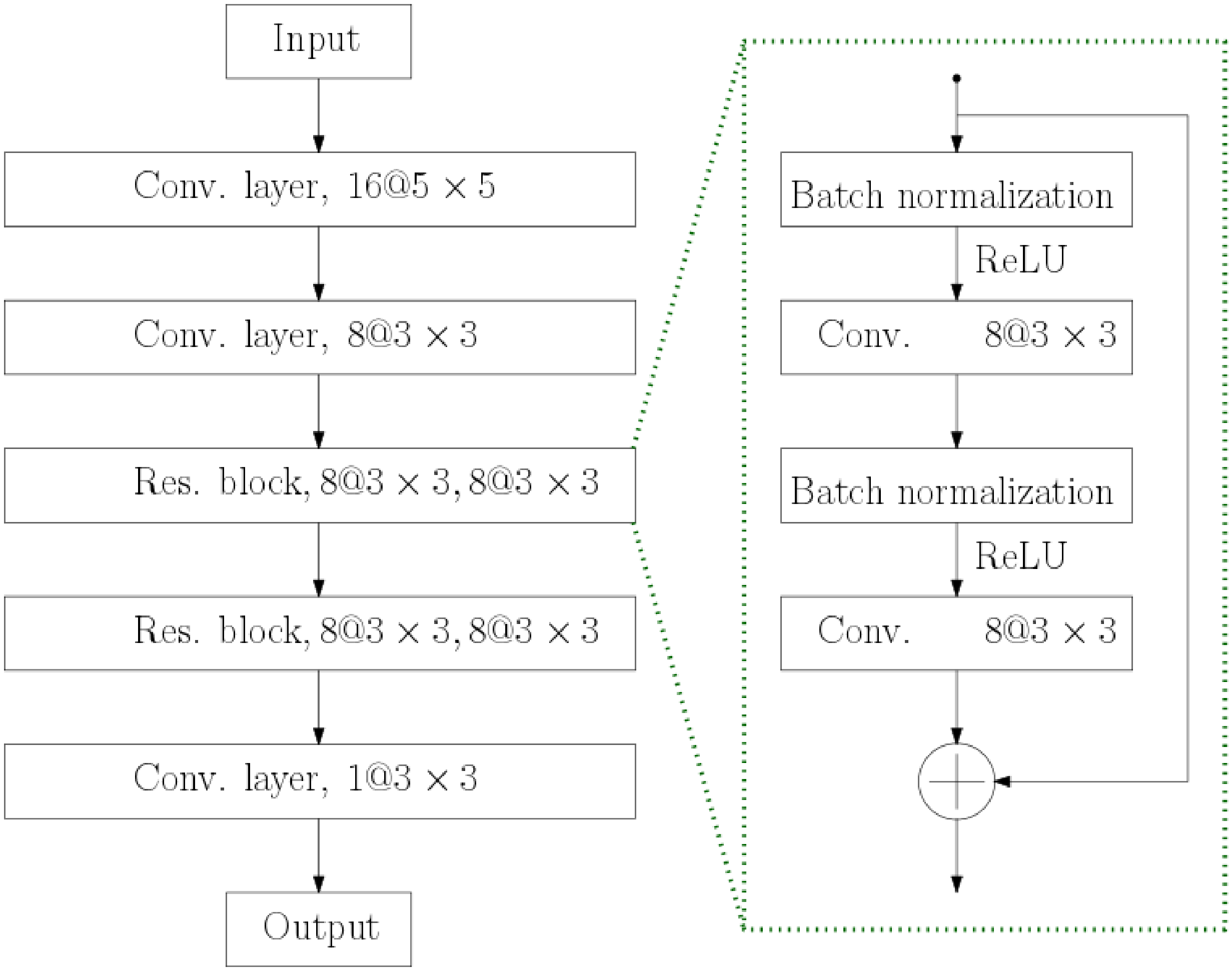


Figure 3: ResNet architecture used in this work. Conv. layers are complex convolution layers.

### III. Training loss functions and learned regularization parameters

In this section, we provide the training and validation losses for the training process of the unfolded network and ResNet. Training was performed in two stages. The first stage consisted of 50 training epochs over 2400 simulated movie patches (20 frames each), while the second stage included additional 20 training epochs over 2400 patches from the first rat (20 frames each). For *in-vivo* validation, 100 consecutive frames from the second rat were chosen randomly. MSE was calculated according to (8) in the main paper.

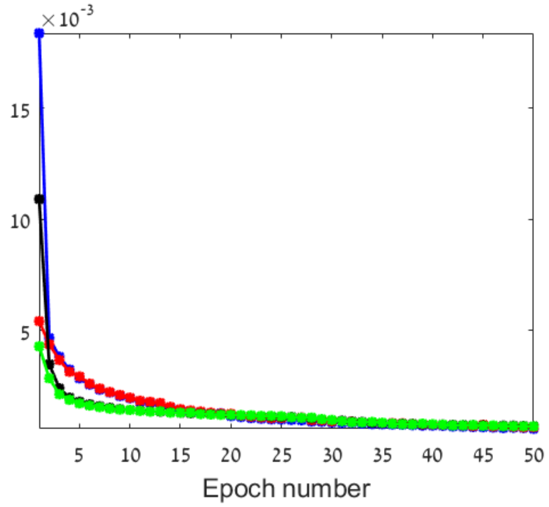


(a) Training on simulation data only.

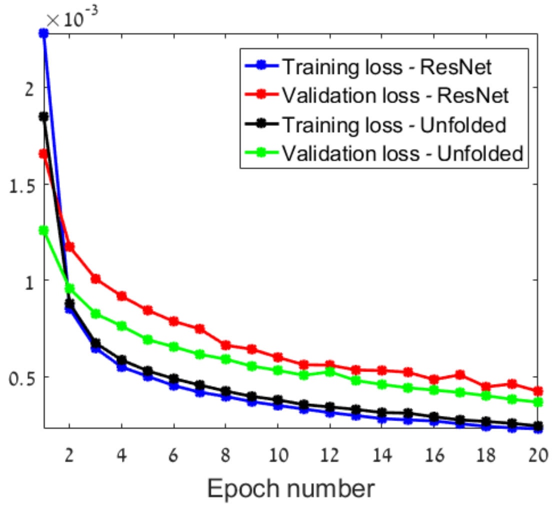


(b) Training on simulation and *in-vivo* data.

Figure 4: Training and validation losses for the unfolded network and ResNet. Left panel shows both training and validation losses for training the unfolded network (10 layers) and ResNet with simulation patches only for 50 epochs. Right panel presents both training and validation losses for the same networks trained with simulation patches for 50 epochs and additional 20 epochs on *in-vivo* data.

Considering Fig. 4, it is evident that when training on simulation data only, the validation curves follow the training loss curves for both networks and are comparable after 20 epochs. This behavior might suggest that the networks over-fit the simulated data, that is, they achieve the best possible recovery for simulated patches. However, in this case, the networks have yet to learn from actual data. In such a case, if the simulation does not represent the data precisely (e.g. different dynamic range, MB concentration, etc.), its performance will degrade when applied to *in-vivo data*, as presented in Section IV. Thus, additional training is performed, as shown in panel (b) of Fig. 4. In this case, the validation losses are higher than the training loss, but now, as presented in the main paper, the networks perform well on *in-vivo* data.

Figure 5 and Fig. 6 illustrate the learned values of $\lambda_{L_i}$ and $\lambda_{S_i}$ for the unfolded network, where $i = 1, \ldots, 10$ indicates the layer number when training on simulation data only and on simulation and *in-vivo* data together.

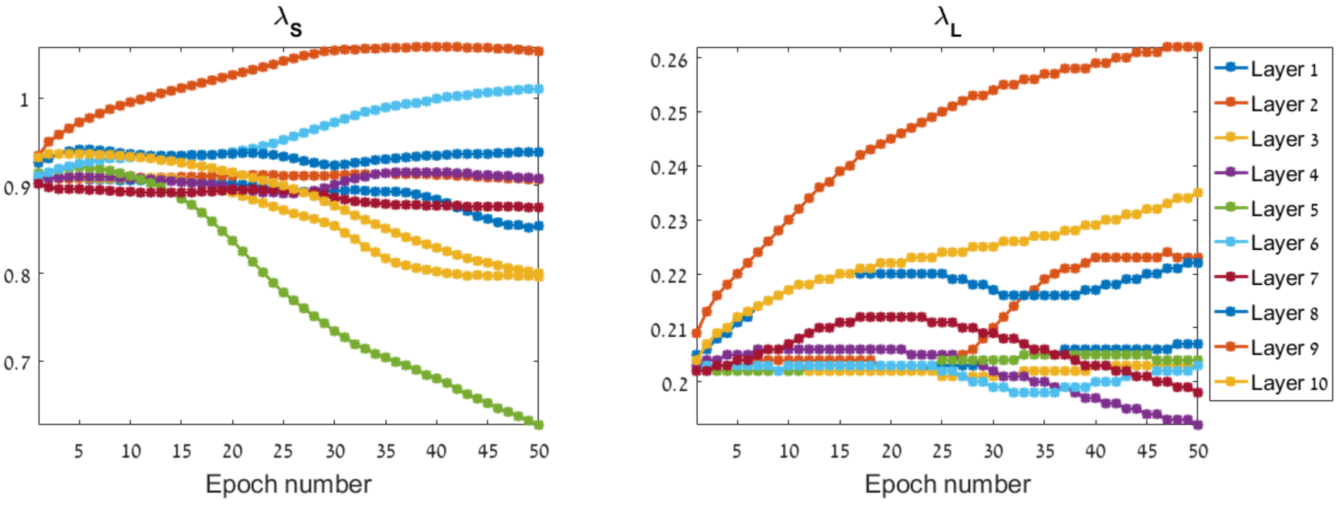


Figure 5: Learned regularization parameters when training on simulation data only.

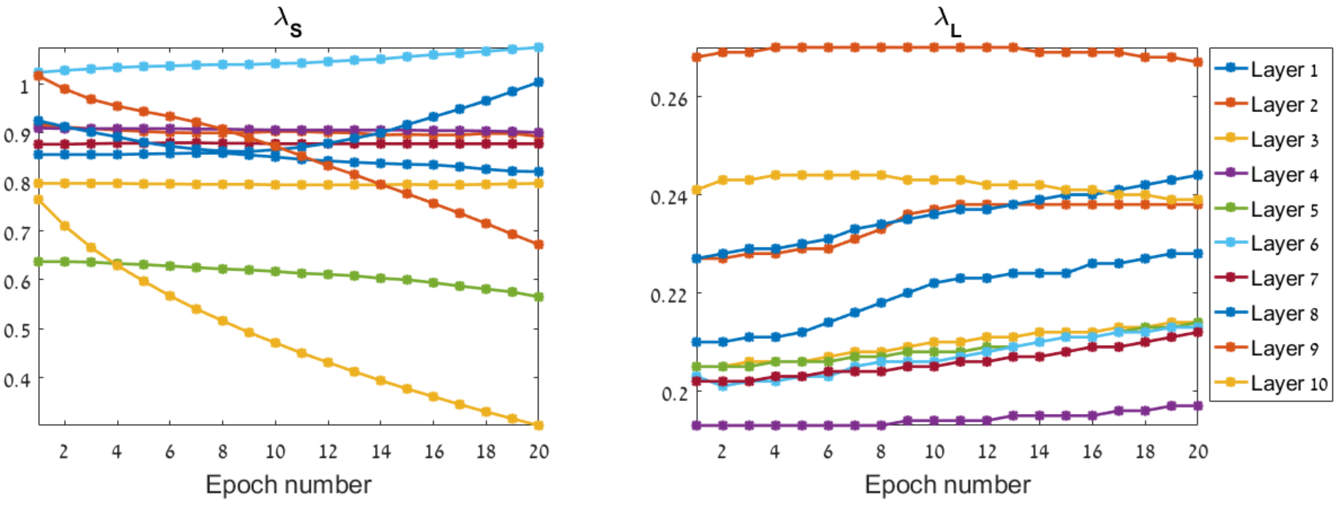


Figure 6: Learned regularization parameters when training on simulation and *in-vivo* data.

Considering both figures, it is evident that most of the regularization parameters do not change considerably when training on *in-vivo* data is performed. As the unfolded network is trained on both simulation and *in-vivo* data, the regularization parameters do not converge to the parameters used in the iterative FISTA algorithm. This also suggests that by performing combined learning on both simulation and experimental data, the network further differs from its iterative counterpart, often leading to improved performance, as presented in the main paper.

### IV. The importance of training on both simulations and in-vivo data

As was described in the main paper and in Section III, the unfolded network outperforms FISTA reconstruction due to the combined training on both simulations and *in-vivo* data. This joint training allows the network to learn both the ”ideal conditions” for MB/tissue separation from the simulations, as well as important features from the experimental data, and achieve robustness to noise and modeling mismatch.

In Fig. 7 we present *in-vivo* results of the network trained in two conditions. Panels (a) and (b) show the output of the network when trained solely on simulated data for 10 epochs and the output when trained on both simulated and experimental data for 10 epochs each, respectively. Panels (c) and (d) show the output of the networks for the same two cases, only now the numbers of training epochs were 50 for simulated data and 20 for *in-vivo* data.

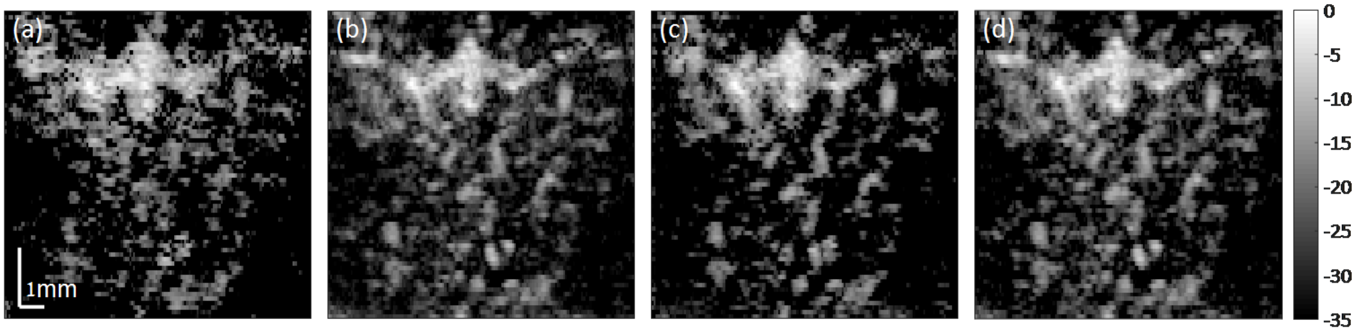


Figure 7: *In-vivo* results when training on simulations only and on simulations and *in-vivo* data together. (a) Training on simulations for 10 epochs. (b) Training on both simulations and experimental data for 10 epochs each. (c) Training on simulations for 50 epochs. (d) Training on both simulations and experimental data for 50 and 20 epochs, respectively. Color bar is in dB.

Considering Fig. 7, clearly when training on a relatively low number of epochs (10), simulated data is not sufficient for good performance on experimental data. On the other hand, when combined with additional 10 epochs of training on *in-vivo* data, the performance of the network improves considerably, and is somewhat similar to the performance of the network result displayed in panel (d). Surprisingly, even when training on simulated data only for enough epochs, in this case 50, the network performs well in recovering the vascular bed of experimental data, as shown in panel (c). However, closer examination shows that albeit the image looks sparser than the image in panel (d), its texture looks more pixel-like than the FISTA and SVD images shown in Fig. 4 of the main paper.

The latter example suggests two things. First, that good results can be obtained by training the network on realistic simulations for enough training epochs. The second is that performance more similar in texture and visual quality to that of non-learning based techniques can be obtained by the combined training on both simulations and experimental data.

### V. Runtime comparison

Here, we compare the run-time performance of both the unfolded network and ResNet, for both training phase and validation phase. Fig. 8 show the time in seconds each network required to train a single epoch and then validate its performance, in yellow, as a function of epoch number. Training was performed for 50 epochs on simulated data. Run-times results for training on *in-vivo* data were similar, and thus are omitted.

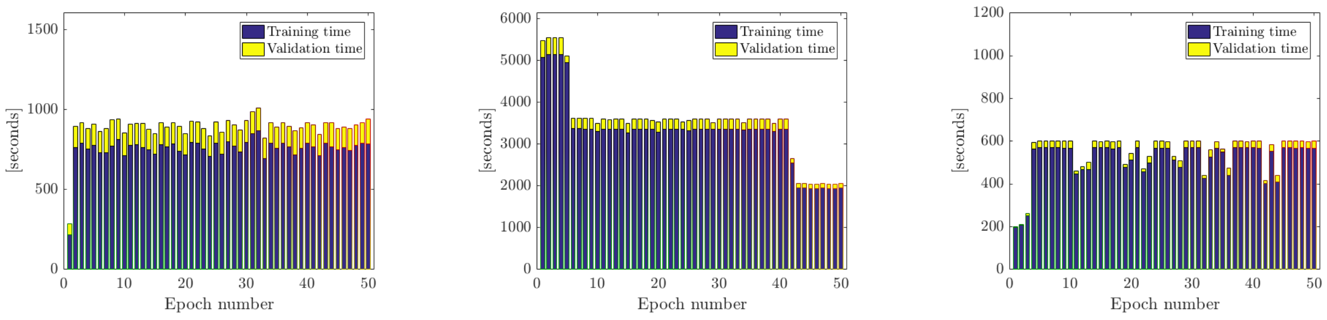


(a) Unfolded network, 1 layer (b) Unfolded network, 10 layers (c) ResNet

Figure 8: Run-time results for training and validation of the unfolded network and ResNet.

Observing Fig. 8, it is evident that the training and validation of the unfolded network is slower compared with ResNet. The 10 layers network is slower by an order of magnitude, but the 1 layer network has a slightly slower runtime. The slower processing and training time of the unfolded network is attributed to the SVD operations required by the network, although faster and more efficient algorithms for SVD computations can be used, as discussed in the Discussion section of the main paper. This figure further supports the conclusions in the main paper. The unfolded network offers a flexible trade-off between execution time and performance, by allowing to choose its depth.

However, the unfolded network has an order of magnitude lower number of trainable parameters and achieves better CNR and CR values, as demonstrated in the main paper. It is also important to remember that the ResNet was not fully trained, rather only its last fully connected layers were trained. This transfer learning process considerably reduces the overall training time.

### VI. Simulation description

As was indicated in the main paper, in this work we increase the number of training examples by training on both experimental and simulated data. In this section, we describe how the simulation was generated. In the simulations we used, pixel size is assumed to be $0.12 \times 0.12\text{mm}^2$ and the number of pixels is $128 \times 128$. Implementation was performed in Python 3.5.2.

#### A. MB signal generation

The overall number of MBs (as well as their initial positions) was generated randomly up to a maximum concentration of $130$ MBs per cm$^{-2}$. MB amplitudes were drawn from a normal complex distribution.

MB velocity magnitudes were generated according to

$$v(x, y, t = 0) = \max(0, v_{\text{det}} \cdot \mathcal{N}(1, 1)),$$

where $v_{\text{det}} = 0.24\text{mm}/dt$, $dt = 0.01\text{s}$ is the imaging frame-rate and $\mathcal{N}(1, 1)$ is a normal distribution with mean 1 and standard deviation 1. MB accelerations were generated according to

$$a_{x/y} = \mathcal{N}(0, \sigma_a),$$

with $\sigma_a = 0.05 \cdot 0.12/dt^2$ and $x/y$ are the lateral and axial directions, respectively.

MB velocity directions are generated in each frame according to

$$v_x^k(t) = v_x^{k-1}(t)\text{cos}(\theta) - v_y^{k-1}(t)\text{sin}(\theta),$$

$$v_y^k(t) = v_x^{k-1}(t)\text{sin}(\theta) + v_y^{k-1}(t)\text{cos}(\theta),$$

with $\theta \sim \text{U}[-30°, 30°]$ and $k$ indicates frame number. MB amplitudes are additionally multiplied by a random factor between 0.9 and 1.1 in each frame.

#### B. Tissue signal generation

To model the tissue signal, we start by generating a sum of five real 2D Gaussian matrices of the same size as the image frames ($128 \times 128$ pixels) with random positions and variances. We then generate a complex random matrix to modulate the envelope of the tissue signal. The real and complex entries are both drawn from a normal distribution with zero mean and standard deviation 1. Both matrices are then multiplied element-wise, and the product is then low-pass filtered (2D real Gaussian matrix of $11 \times 11$ pixels). The resulting signal's envelope, denoted as $\mathbf{B} \in \mathcal{R}^{I \times J}$ mimics the texture of the tissue signal. Thus, the overall pixels' values are random, but locally they are correlated.

The next step involves the generation of a phase matrix, same size as before. It's entries are drawn from a Gaussian distribution in the following manner

$$\boldsymbol{\theta} \sim \mathcal{N}(\alpha, \sigma_\theta),$$

with a mean drawn from a uniform distribution in the range $\alpha \sim [0°, 180°]$ and standard deviation of $\sigma_\theta = 15°$. The resulting complex tissue signal is given by

$$\mathbf{T}[i, j] = \mathbf{B}[i, j]e^{j\boldsymbol{\theta}[i,j]},\ i, j \in [I, J].$$

The next stage in generating the tissue signal is to apply spatial deformations in each frame, to mimic tissue movement during the acquisition period. To this end we start by generating four different $4 \times 4$ kernels, denoted as flow filters. The entries of those kernels, are positive and their sum equals to one. For each new frame, we generate additional four filters, with entries drawn from a Gaussian distribution with zero mean and standard deviation of 0.1. For each flow filter we add the corresponding new filter and for each pixel we take the maximum between the latter value and 0.1. The resulting new filter is normalized such that all entries sum to one and the entries are non-negative.

Once the flow filters have been updated for the current frame, they are convolved with $\mathbf{T}$, to get four different images. We then divide each of the four images into $4 \times 4$ blocks. The final image is generated by dividing an empty matrix into $4 \times 4$ blocks, and for each block choosing randomly one of the corresponding blocks from the four images. This process ensures that blocks in the same neighborhood share the same movement pattern, but in the whole image, the pattern is random.

#### C. Simulation of the PSF

Once the (complex) MB frame and tissue frame are generated and summed with complex Gaussian noise, the resulting frame is convolved with the PSF. The PSF is modeled as a 2D real Gaussian kernel with standard deviations of $0.14$mm in the lateral and $0.32$mm axial dimensions, taken from the *in-vivo* data.
